# Vγ1^+^ γδ T cell-derived IL-4 initiates CD8 T cell immunity

**DOI:** 10.1101/2024.12.06.627130

**Authors:** Shirley Le, Declan Murphy, Shangyi Liu, Zhengyu Ge, Rose May, Anton Cozijnsen, Thomas Burn, Charlie Jennison, Annabell Bachem, Calvin Xu, Hui-Fern Koay, Jan Schröder, Stephanie Gras, Ian A. Cockburn, Sammy Bedoui, Laura K. Mackay, Geoffrey I McFadden, Daniel Fernandez-Ruiz, William R Heath, Lynette Beattie

**Affiliations:** Department of Microbiology and Immunology and The Peter Doherty Institute for Infection and Immunity, University of Melbourne, Parkville, Australia; School of Biosciences, University of Melbourne, Parkville, Australia; Computational Sciences Initiative, Department of Microbiology and Immunology and The Peter Doherty Institute for Infection and Immunity, University of Melbourne, Parkville, Australia; Infection and Immunity program, La Trobe Institute for Molecular Science (LIMS), La Trobe University, Bundoora, Australia; Department of Biochemistry and Chemistry, School of Agriculture, Biomedicine and environment, La Trobe University, Bundoora, Australia; Department of Biochemistry and Molecular Biology, Monash University, Clayton, Australia; Division of Immunology and Infectious Disease, John Curtin School of Medical Research, Australian National University, Canberra, Australia; School of Biomedical Sciences, Faculty of Medicine & Health, and the UNSW RNA Institute, The University of New South Wales, Kensington, NSW 2052, Australia

## Abstract

Dendritic cells (DC) are pivotal for initiating adaptive immunity, a process triggered by the activation of DC via pathogen products or damage. Here, we describe an additional layer to this process, essential when pathogen-derived signals alone cannot directly achieve full DC activation. Immunisation with sporozoites from *Plasmodium* leads to CD8 T cell priming in a complex response that is initiated by a collaboration between conventional type 1 DC (cDC1) and γδ T cells. We unveil a pivotal initiating role for Vγ1^+^ γδ T cells, as they directly supply IL-4 to DC and CD8 T cells. IL-4 synergises with a CD4 T cell-derived CD40L signal to induce IL-12 production by cDC1. Both IL-12 and IL-4 then directly signal CD8 T cells, with synergy between these cytokines driving enhanced IL-12 receptor expression and expansion of responding CD8 T cells. This study reveals a key role for Vγl^+^ γδ T cells in initiating CD8 T cell immunity to *Plasmodium*. More broadly, it shows that responses to some pathogens require help from innate-like T cells to pass an initiation threshold and further amplify the response in a process underscored by IL-4 production.

## Introduction

Priming of CD8 T cells is a complex process that relies on antigen presentation by DC that are sufficiently activated to provide all the necessary costimulatory signals. Classically, DC activation involves pattern recognition receptor (PRR) based detection of the pathogen itself, and inflammatory cues associated with the encounter. These signals increase costimulatory molecule expression on the DC and enhance MHC-II expression for antigen presentation to CD4 T cells^1^. The CD4 T cells then provide CD40L, thereby licensing the DC^2–4^ and stimulating production of cytokines that contribute to T cell differentiation and expansion^5^. DC that are mature but not completely activated (or ‘immunogenic’) fail to produce pro-inflammatory cytokines^1^, resulting in impaired T cell-mediated immunity^6^. Immune responses to pathogens that induce strong pattern recognition receptor (PRR) signalling in DC are less dependent upon CD40L-mediated help from CD4 T cells^7^. Conversely, the responses to some pathogens^7^ and other poor immunogens, are more dependent upon CD4 T cell help for full DC activation and licensing^8^. CD8 T cell responses to radiation attenuated *Plasmodium* sporozoites (RAS), are completely dependent upon help from CD4 T cells^9,10^, placing *Plasmodium* sporozoites in the category of poor immunogens.

The cytokines produced in response to DC activation and classically associated with differentiation and expansion of CD8 T cells include type I interferons, IL-2, IL-12 and IFNγ^11^, but not commonly IL-4. Regarded as a prototypical Th2 cytokine, IL-4 has well-defined roles in CD4 T cells^12^ but recent studies have suggested that IL-4 may also have potent and context dependent roles in CD8 T cells, that have implications for T cell-based immunotherapy success^13,14^ or failure^15^.

One model known to be dependent on IL-4 is intravenous immunisation with RAS^16,17^. RAS injection induces protective immunity due to activation of CD8 T cells in the spleen by cDC1^18^. These activated CD8 T cells then expand and a proportion of them differentiate into memory phenotype cells including liver resident memory T cells (T_RM_) that can protect against re-infection^19^. Strikingly, in both animal models and human vaccination, γδ T cells are associated with the success of this response^20^, though just how these cells contribute is unclear.

γδ T cells are innate sensors^21^ that respond to infection, stress, or damage^21,22^. Once activated, γδ T cells exert diverse effector functions including the rapid release of cytokines^23–25^. In mice, Vγ1^+^, and Vγ4^+^ populations^26^ are found in lymphoid tissues including the spleen^22^. In humans, the predominant γδ T cell subsets express either Vδ1^+^, Vδ2^+^ or Vδ3^+^, with the majority of Vδ2^+^ cells preferentially pairing with Vγ9^27^. Vγ9^+^Vδ2^+^ T cells are the most abundant γδ T cell population in human peripheral blood and are also found in the spleen^28^. Strong correlations between γδ T cell functions and the generation of protective immunity following RAS vaccination have been established in humans and in mice^20,29,30^. However, the molecular pathways driving these associations, the specific γδ T cell populations involved, and the molecular signals they contribute to protective immunity remain unclear.

Here, we show that in mice the Vγ1^+^ subset of γδ T cells initiate the CD8 T cell response to liver stage malaria parasites via IL-4 production. Our data have implications for our understanding of how CD8 T cell responses to poor immunogens can be strengthened early in the priming phase and reveal a role for Vγ1^+^ γδ T cells and IL-4 in enhancing CD8 T cell responses.

## Results

### γδ T cells affect early expansion of CD8 and CD4 T cells

The T cell response to intravenously injected RAS is initiated largely in the spleen with priming and expansion occurring in this site, followed by recirculation and resultant rapid accumulation of activated T cells in the liver^18^. A proportion of these cells will then differentiate into memory T cell subsets, including protective liver T_RM_ cells^19^. To understand if the previously described role for γδ T cells in the generation of memory T cell responses in RAS vaccinated mice^20^ was due to a role for γδ T cells in T cell priming, or in the generation of memory T cells, we used PbT-I TCR transgenic T cells to study the response at an antigen-specific level. These T cells recognise an H-2K^b^-restricted epitope (PbRPL6_120-127_) from the *P. berghei*-derived RPL6 protein^31,18,32^. PbT-I cells were adoptively transferred into B6 or δ^−/-^ mice and responses assessed 6 days after vaccination with RAS (**Figure 1a, b**). This revealed a defect in the accumulation of PbT-I cells in the spleen and consequently, the liver of δ^−/-^ mice. As a result of this early failure to respond to RAS, fewer memory PbT-I cells were found in either organ after 3 weeks (**Figure 1c, d**), with impaired formation of memory T cells subsets, including liver T_RM_ cells (**Figure 1e and f**).

**Figure 1.**
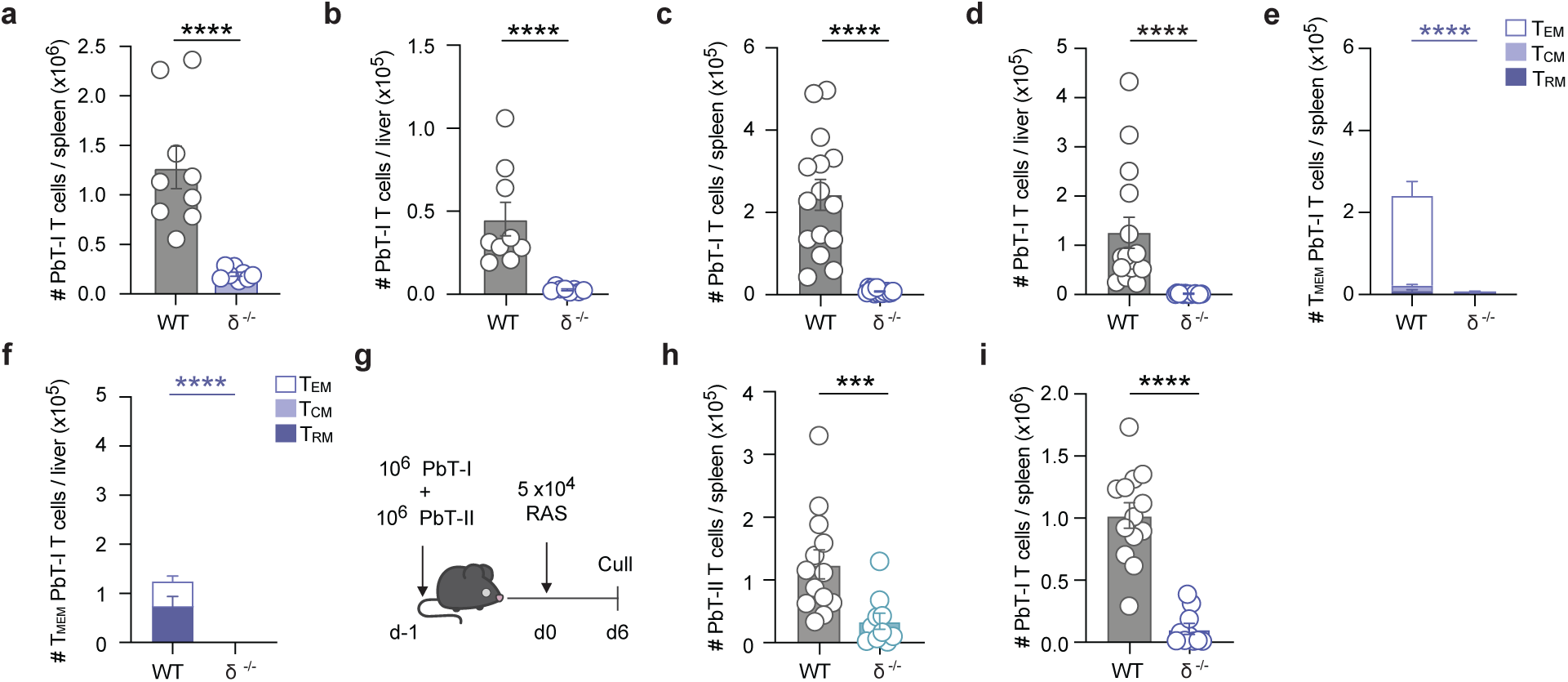
γδ T cells are essential for liver T_RM_ cell generation. 10^6^ RPL6-specific transgenic CD8^+^ T cells (PbT-I) were transferred into mice one day prior to vaccination with 5 ×10^4^ RAS. PbT-I cells were analysed either 6 or 23 days later. (**a and b)** Enumeration of PbT-I cells in the (**a**) spleen or (**b**) liver of B6 or *Tcrd*^−/-^ mice, 6 days after immunisation. (**c and d)** Numbers of PbT-I cells in the (**c)** spleen or (**d**) liver of B6 or *Tcrd*^−/-^ mice, 23 days after vaccination. **(e and f**) Quantified memory T cell (T_MEM_) subsets; central memory T_CM_ (CD62L^+^, CD69^−^), effector memory T_EM_ (CD62L^−^, CD69^−^) and resident memory T_RM_ (CD62L^−^, CD69^+^) within the CD44+ PbT-I cell compartment of the (**e**) spleen or (**f**) liver at day 23. **(g)** Experimental design for (**h and i**). 10^6^ transgenic CD8^+^ (PbT-I) and CD4^+^ (PbT-II) T cells were transferred into mice one day prior to vaccination with 5×10^4^ RAS **(h)** PbT-II or (**i**) PbT-I cell counts in the spleen on day 6. Data pooled from (**a-b**) 2 or (**c-i**) 3 independent experiments with n = 9-15 mice per group. Error bars indicate mean + SEM. Data were log-transformed and compared using an unpaired t test. ****p<0.0001, ***p<0.001, **p<0.01, *p<0.05, ns=p>0.05.

CD8 T cells within the endogenous repertoire specific for PbRPL6_120-127_^31^ also showed impaired accumulation in the spleen and liver of δ^−/-^ mice 6 days after RAS vaccination (**Supplementary Figure 1a-c)** and, analogous to the PbT-I cells, did not form memory in the spleen or liver (**Supplementary 1d-g**). The response of PbT-I cells therefore phenocopied the response of *Plasmodium*-specific T cells within the endogenous repertoire. Together these findings showed that γδ T cells were required for the initiation phase of the CD8 response to RAS, explaining why memory responses were blunted.

Of note, the poor initial response in δ^−/-^ mice only extended to vaccination with the sporozoite form of this parasite as responses to blood-stage parasites (irradiated infected red blood cells) were unaffected (**Supplementary Figure 1h**), despite this response being categorised as a relatively weak response based on its dependence on CD4 T cell help^33^.

We next assessed whether the initiation of the CD4 T cell response to RAS vaccination was also γδ T cell dependent. Here we utilized PbT-II cells^33^ from the MHC-II restricted P*. berghei*-specific TCR transgenic line^34^. Adoptive transfer of both PbT-I and PbT-II cells into B6 or δ^−/-^ mice (**Figure 1g**) showed that splenic accumulation of PbT-II cells was dependent upon γδ T cell-mediated help in response to RAS vaccination (**Figure 1h**). As a result, fewer PbT-II cells were present in the livers of δ^−/-^ mice (**Supplementary Figure 1i**). PbT-I cell accumulation in the spleen (**Figure 1i**) and liver (**Supplementary Figure 1j**) of the δ^−/-^ mice was also lower than the controls, demonstrating that the addition of large numbers of naïve antigen-specific CD4 T cells could not rescue the response of CD8 PbT-I cells when γδ T cells were absent.

### Vγ1^+^ γδ T cells initiate immunity to RAS

To ascertain when γδ T cells affect CD4 and CD8 T cell responses, we utilised the pan-TCRδ blocking antibody (α-γδ, clone GL3)^35^ to block γδ T cell function at the time of, or 24 hr after RAS vaccination (**Figure 2a**). Blocking γδ T cell function prior to RAS vaccination impaired the PbT-I response comparably to that previously observed in δ^−/-^ mice (**Figure 2b**). In contrast, blocking γδ T cell function from 24 hr post-RAS injection had a milder effect (**Figure 2b**). These data suggest that the first 24 hr are the crucial window of γδ T cell activation.

**Figure 2.**
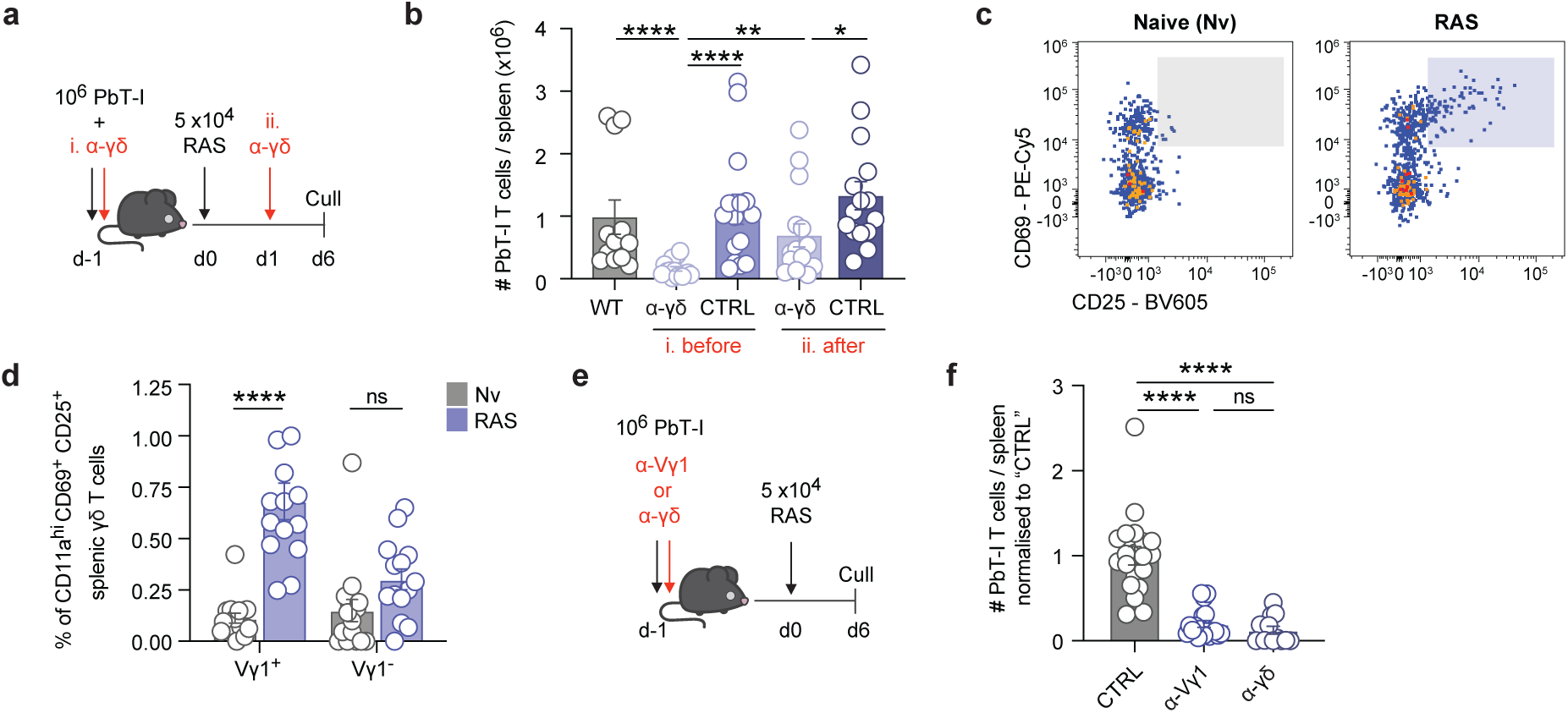
Splenic Vγ1^+^ γδ T cells initiate immunity to RAS. **(a)** Experimental design for (**b**). 10^6^ PbT-I cells were transferred to recipient mice one day prior to RAS vaccination. Mice were treated with an α-pan-γδ-TCR (α-γδ, clone GL3) or an isotype control Ab, “CTRL”, either (i) before RAS, or (ii) 24 hours after RAS. **(b)** Numbers of PbT-I cells in the spleen were assessed on day 6 post-vaccination. **(c)** Representative flow cytometry plots of Vγ1^+^ CD11a^hi^ γδ T cells in naïve (left) or RAS-vaccinated (right) TCRδ-GDL mice 24 hours after injection. **(d)** Frequency of activated (CD69^+^ CD11a^hi^ CD25^+^) Vγ1^+^ or Vγ1^−^γδ T cells in naïve or RAS-vaccinated TCRδ-GDL mice 24 hours post-injection. **(e)** Experimental design for (**f**) **(f)** Normalised splenic PbT-I cell numbers 6 days post-vaccination in isotype control Ab treated “CTRL”, α-Vγ1-treated (clone 2.11) or α-pan-γδ-TCR treated (clone GL3) mice. Numbers were normalised by expressing them as a fraction of the mean of the “CTRL” group. Data pooled from (**a-d**) 3 or (**e, f**) 4 independent experiments with n= 9-20mice per group. Error bars indicate mean + SEM. Data were log-transformed and compared using an (**b, f**) ordinary one-way or (**d**) two-way ANOVA. ****p<0.0001, ***p<0.001, **p<0.01, *p<0.05, ns=p>0.05.

Knowing that the first 24 hours were critical, we next examined early γδ T cell activation. Splenic γδ T cells were split into the two major populations found in murine spleen, Vγ1^+^ or Vγ1^−^ populations (**Supplementary Figure 2a**). Up-regulation of the canonical T cell activation markers CD69, CD25 and CD11a in the Vγ1^+^, but not the Vγ1^−^ γδ T cell population, indicated that Vγ1^+^γδ T cells were preferentially activated by RAS vaccination (**Figure 2c, d and Supplementary Figure 2b**). Blocking the Vγ1^+^ γδ T cell population via injection of a Vγ1 blocking antibody (clone 2.11)^36^ (**Figure 2e, f**) had the same effect on PbT-I accumulation as blockade of the entire γδ T cell population (α-γδ). Vγ1^+^ γδ T cells therefore initiate the CD8 T cell response to RAS in mice.

**Antigen presentation of *Plasmodium*-derived peptides is intact in the absence of**γδ **T cells**

Due to the very early timing of γδ T cell activation, and the effect of Vγ1^+^ γδ T cells on CD8 T cell accumulation in the spleen, we hypothesised that the Vγ1^+^ γδ T cells may impact the ability of cDC1 to present antigen to CD8 T cells in response to RAS vaccination. To investigate this possibility, we examined initial up-regulation of CD69 within OT-I T cells following injection of SIINFEKL expressing CS5M sporozoites (CS5M-RAS). OT-I T cells recapitulated the phenotype of PbT-I cells in δ^−/-^ mice, with impaired responses in the spleen and liver 6 days after vaccination with CS5M-RAS (**Supplementary Figure 3a and b**). At early time points post-CS5M-RAS vaccination, equal proportions (**Figure 3a and b**) and numbers (**Figure 3c**) of cell trace violet (CTV) labelled OT-I T cells up-regulated cell surface expression of CD69 in the spleens of B6 and δ^−/-^ mice. These data indicated that responding T cells had access to antigenic signals capable of up-regulating CD69 and initiating some proliferation even when γδ T cells were absent. Failure to accumulate on day 6 (**Supplementary Figure 3a and b**), however, suggested a lack of signals for extended T cell expansion, differentiation and/or survival. This conclusion strongly suggested a lack of co-stimulation or cytokines in the T cell priming process, raising the intriguing possibility that γδ T cells are crucial for triggering upregulation of co-stimulation-like signals on cDC1.

**Figure 3.**
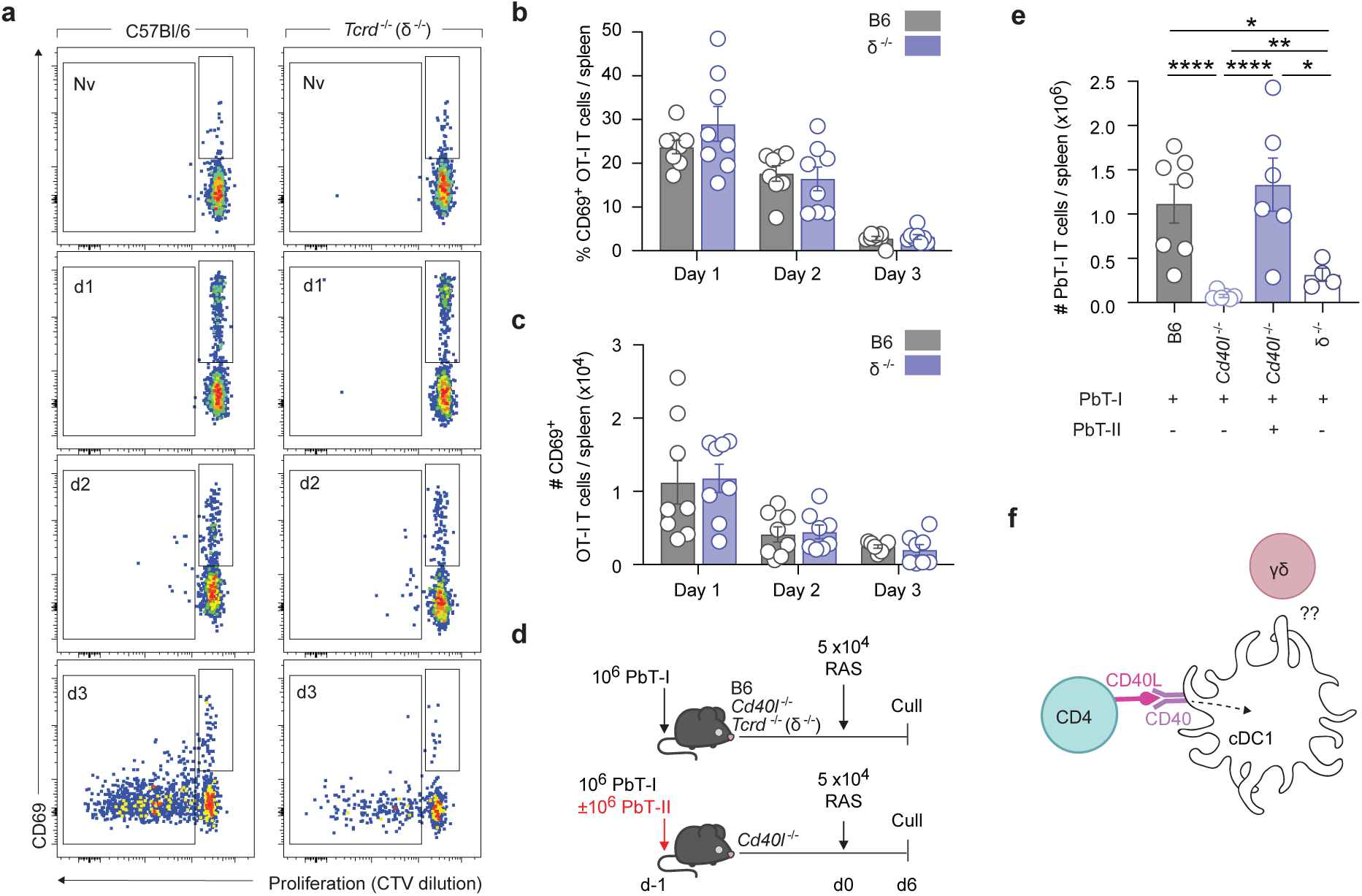
Antigen presentation and CD40-CD40L signalling is intact in the absence of γδ T cells. 10^6^ CTV-labelled OVA-specific CD8^+^ (OT-I) T cells were transferred into B6 or *Tcrd*^−/-^ recipient mice, which were subsequently vaccinated one day later with 5 ×10^4^ CS5M-RAS. OT-I cell numbers were assessed on day 1, day 2 and day 3 after transfer. **(a)** Representative flow cytometry plots of CD44^+^ OT-I cells in the spleen at the indicated time points. (**b, c**) (**b**) Frequency and (**c**) number of CD69^+^ CTV^hi^ OT-I cells in the spleen of either B6 or *Tcrd*^−/-^ mice. **(d)** Experimental design for (**e**). B6, *Tcrd*^−/-^ and *Cd40l*(*Cd154*)^−/-^ mice received PbT-I cells with or without PbT-II cells co-transferred one day prior to vaccination with 5×10^4^ RAS. **(e)** PbT-I cell counts in the spleen at day 6 post-vaccination. **(f)** CD40 signalling to DC is intact when CD40L sufficient T cells are provided, showing that CD4 T cells, not γδ T cells provide CD40L in this system. Data pooled from 2 independent experiments with (**a-d**) n = 8 or (**f**) n= 6 mice. Error bars indicate mean + SEM. Data were log-transformed data and compared using an unpaired t test **(d)** or an ordinary one-way ANOVA (**f**). ****p<0.0001, ***p<0.001, **p<0.01, *p<0.05, ns=p>0.05.

### γδ T cells are not required to supply the CD40L signal to cDC1

As a deficiency in co-stimulation was implicated, we hypothesised that γδ T cells may provide CD40L for signalling CD40 on cDC1, a signal known to be essential in the response to RAS^37^. To investigate this possibility, PbT-I cells were transferred into WT, δ^−/-^ or *Cd40l^−/-^* mice. CD8 T cell expansion is dependent on help from CD4 T cells in the RAS model^9,10^, and these CD4 T cells were previously presumed to provide the CD40L signal. To test this assumption, one group of *Cd40l^−/-^* mice were also given CD40L sufficient PbT-II cells (**Figure 3d**). PbT-I cell accumulation in the spleen 6 days after RAS injection was severely diminished in *Cd40l^−/-^* mice (**Figure 3e**), confirming that CD8 T cell accumulation in this model is dependent on CD40L signalling. Addition of CD40L sufficient PbT-II cells, however, rescued the PbT-I cell response, even though γδ T cells lacked expression of CD40L in these mice (**Figure 3e**). Therefore, CD4 T cells can provide the CD40L signal required for the response to RAS suggesting that γδ T cells are not required to provide this co-stimulatory function. Furthermore, rescue by CD40L sufficient PbT-II cells showed that this signal is normally provided by CD4 T cells but is insufficient to ensure an appropriate CD8 T cell response if γδ T cells are absent (**Figure 3f**).

### IL-4 is required for the CD8 T cell response to RAS vaccination

As CD40L was not the crucial signal provided by γδ T cells, we rationalised that they may provide ‘signal 3’ i.e. a cytokine signal, for differentiation and expansion of responding T cells. Previous studies suggested IL-4 is a crucial factor in the generation of memory CD8 T cells in the liver following RAS vaccination^16,38^. Given that splenic Vγ1^+^ γδ T cells have been reported to secrete IL-4^24^, we next investigated the role of IL-4 in the PbT-I cell response either via transfer into *Il4*^−/-^ hosts (**Figure 4a and b**) or by antibody-mediated blockade (**Supplementary Figure 4a and b**). PbT-I cells did not accumulate in the spleen or liver in the absence of IL-4, showing that IL-4 was crucial, and the absence of IL-4 mirrors the effect of γδ T cell deficiency.

**Figure 4.**
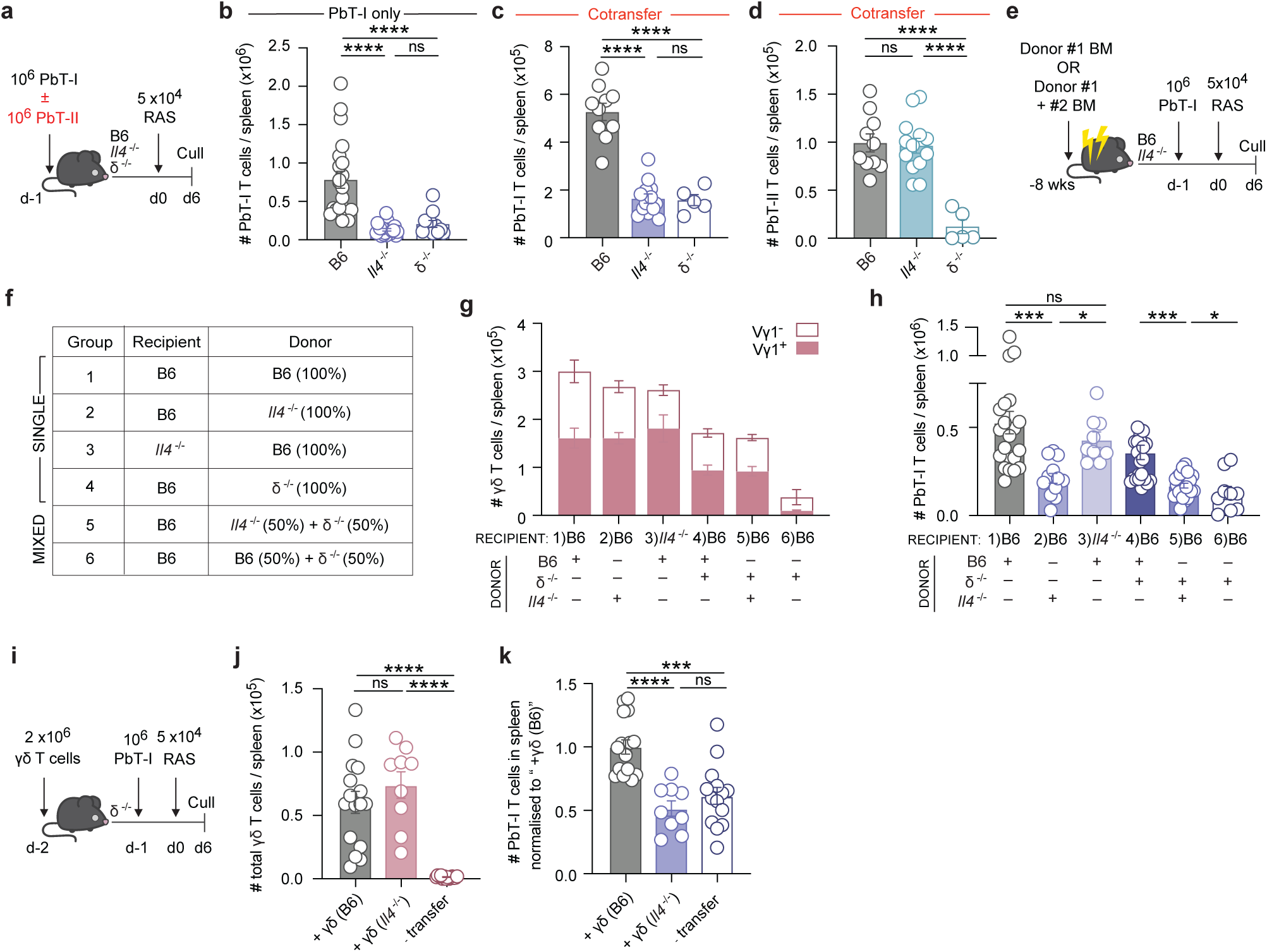
IL-4 is required for the CD8 T cell response to RAS vaccination. 10^6^ PbT-I cells were transferred into B6, *Il4*^−/-^ or *Tcrd*^−/-^ mice one day prior to immunisation. An additional cohort of *Il4*^−/-^ mice received both 10^6^ PbT-I cells and 10^6^ PbT-II cells. **(a)** Experimental design for (**b-d**). **(b)** Number of PbT-I cells in the spleen of B6, *Il4*^−/-^ or *Tcrd*^−/-^ mice at 6 days post-vaccination. (**c and d**) Numbers of (**c**) PbT-I cells and (**d**) PbT-II cells in the spleen of B6, *Il4*^−/-^ or *Tcrd*^−/-^mice that received both PbT-I and PbT-II cells. **(e)** Experimental design for (**f-h**). Chimeras were given 10^6^ PbT-I cells one day prior to RAS vaccination then analysed 6 days later. **(f)** Single and mixed bone marrow chimeras were prepared using bone marrow from B6, *Tcrd*^−/-^, *Il4*^−/-^ donors as indicated. **(g)** Enumerated splenic γδ T cells in single and mixed chimeras separated by Vγ1 expression. **(h)** Splenic PbT-I cell numbers were assessed in chimeras, 6 days after RAS vaccination. **(i)** Experimental design for (**j and k**). *Tcrd*^−/-^ mice received γδ T cells enriched from the spleens of either B6 or *Il4*^−/-^ donors one day prior to transfer of PbT-I cells. Recipient mice were subsequently vaccinated with RAS and analysed 6 days later. **(j)** Quantified splenic γδ T cells 6 days after RAS vaccination in *Tcrd*^−/-^ mice that received γδ T cells from B6 (+γδ B6), or *Il4*^−/-^ mice (+γδ *Il4^−^*) or no γδ T cells (no transfer). **(k)** Normalised PbT-I cell counts in the spleen 6 days after RAS vaccination. Counts were normalised by expressing them as a fraction of the mean of the “+ γδ (B6)” group. Data pooled from **(b, g and h**) 4 or **(c, d, j and k**) 3 independent experiments where n=5-23 mice per group. Error bars indicate mean + SEM. Data in (**b-d, g, h, j, k**) were log-transformed and compared using an ordinary one-way ANOVA. ****p<0.0001, ***p<0.001, **p<0.01, *p<0.05, ns=p>0.05

To further investigate if, as previously reported, CD4 T cells are the relevant source of IL-4^16^, we transferred IL-4 sufficient PbT-I cells with, or without, IL-4 sufficient PbT-II cells into B6 or *Il4*^−/-^ mice and measured T cell accumulation as the output (**Figure 4a**). Accumulation of PbT-I cells was impaired in the *Il4*^−/-^ mice even in the presence of antigen-specific CD4 T cells that could make IL-4 (**Figure 4c**). In contrast, antigen-specific CD4 PbT-II cells expanded in the spleen in both B6 and *Il4*^−/-^ mice (**Figure 4d**) demonstrating two phenomena; 1) the CD4 T cell response was not IL-4 dependent and 2) IL-4 sufficient CD4 T cells could not rescue the antigen-specific CD8 T cell response. CD4 T cells, therefore, are not the crucial source of IL-4 for CD8 T cell accumulation following RAS vaccination.

### γδ T cells produce IL-4 in experimental malaria

To determine if γδ T cells provide IL-4 in response to RAS vaccination in mice, we generated several groups of mixed bone marrow (BM) chimeras (**Figure 4e, f**). Eight weeks after reconstitution, the γδ T cell compartment was reconstituted as expected (**Figure 4g**). Assessment of the PbT-I cell response 6 days after RAS vaccination demonstrated impaired PbT-I cell accumulation in the spleens of *Il4*^−/-^+δ^−/-^→B6 chimeras (group 5) when compared to B6+δ^−/-^→B6 chimeras (group 4) (**Figure 4h**). These data demonstrated that γδ T cells were the source of IL-4, since B6+δ^−/-^→B6 chimeras contained γδ T cells that could produce IL-4, whereas the *Il4*^−/-^+δ^−/-^→B6 chimeras lacked these cells. These data strongly suggest that γδ T cells are the essential source of IL-4 for the initiation of an effective CD8 T cell response to RAS vaccination.

To further confirm that γδ T cells are the crucial source of IL-4, we reconstituted δ^−/-^ mice with splenic γδ T cells from either WT *or Il4*^−/-^ donors and tested whether these cells could rescue the PbT-I cell accumulation following RAS vaccination. TCRδ-deficient mice received either splenic γδ T cells or no γδ T cell transfer (**Figure 4i**). Six days after RAS vaccination (8 days after γδ T cell transfer), γδ T cells were recovered from the spleens demonstrating that the transfer was effective (**Figure 4j**). As hypothesised, B6-derived but not *Il4*^−/-^-derived γδ T cells supported PbT-I cell accumulation in the spleen in response to RAS vaccination, confirming that these cells provide IL-4 in this vaccination setting (**Figure 4k**).

### IL-4 has a dual mechanism of action by signalling both CD8 T cells and cDC1 to initiate the CD8 response to RAS

IL-4 is a prototypical T helper 2 cytokine that has potent effects on CD4 T cell differentiation and function^12^, but less well studied impacts on CD8 T cells. We therefore asked if the RAS-induced γδ T cell-derived IL-4 acted directly on the responding CD8 and/or CD4 T cells. To address this question, we used CRISPR-Cas9 to delete the IL-4Rα gene (*Il4ra*) in T cells. Purified naïve PbT-I and/or PbT-II cells had *Il4ra* (or *Cd19* as a control) ablated prior to transfer and immunisation with RAS (**Figure 5a and Supplementary Figure 5a)**. Comparison of the number of PbT-I cells six days after vaccination showed that direct IL-4Rα signalling in PbT-I cells was important for PbT-I cell accumulation in the spleen (**Figure 5b**). These data show that in the context of RAS vaccination direct IL-4 signals to CD8 T cells affect their expansion.

**Figure 5.**
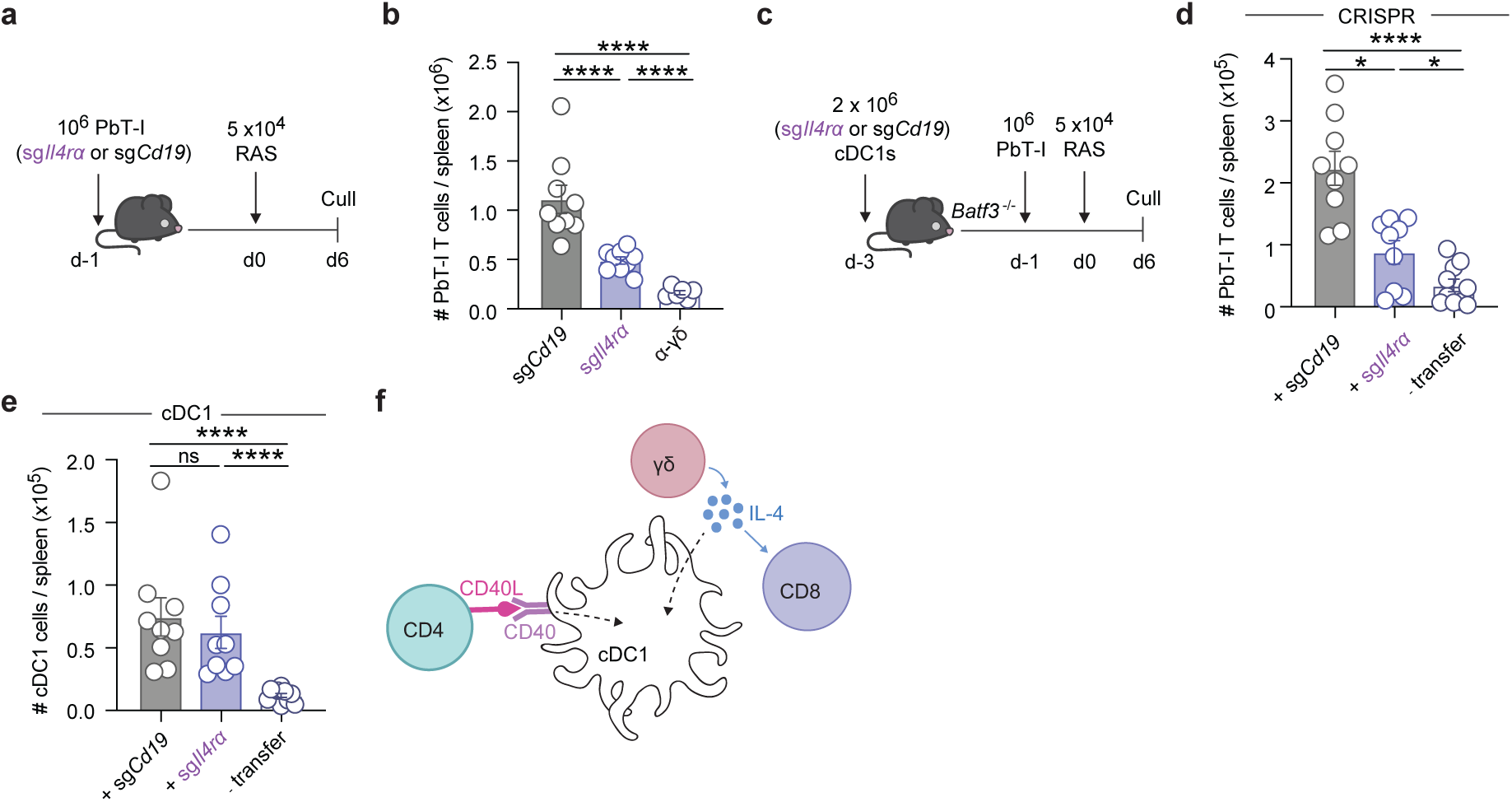
CD8 T cells and dendritic cells respond directly to IL-4. **(a)** Experimental design for (**b**). 10^6^ CRISPR-Cas9 ablated sg*Cd19* (control) or sg*Il4ra* CD8^+^ PbT-I cells were transferred into recipients one day prior to vaccination with RAS.(**b**) Numbers of PbT-I cells in the spleen on day 6 post-vaccination. **(c)** Experimental design for (**d, e**). 2×10^6^ B16.Flt3L-elicted sg*Cd19* (control) or sg*Il4ra* CRISPR/Cas9-edited CD24+ cDC1 were transferred into *Batf3*^−/-^ recipients then given 10^6^ CD8^+^ PbT-I cells two days later. Recipient mice were vaccinated with 5×10^4^ RAS and analysed on day 6 post-RAS. (**f**) Number of PbT-I cells in the spleens of *Batf3*^−/-^ mice that received CRISPR-Cas9 edited cDC1s, or no cDC1s (- transfer). **(d)** Splenic cDC1 counts in *Batf3*^−/-^ mice that received CRISPR-Cas9 edited cDC1s (+sg*Cd19* or +sg*Il4ra*), or no cDC1s (- transfer). (**g**) IL-4 acts directly on CD8 T cells and cDC1s for CD8 T cell accumulation. Data pooled from 2 independent experiments where n=6-16 mice per group. Error bars indicate mean + SEM. Data were log-transformed and compared using an ordinary one-way ANOVA. ****p<0.0001, ***p<0.001, **p<0.01, *p<0.05, ns=p>0.05

PbT-I cells lacking the IL-4R still retained a modest capability to accumulate when compared to a complete failure in the absence of γδ T cells (**Figure 1a**), or IL-4 itself (**Figure 4b**). This suggested that IL-4 has an additional activity that contributes to CD8 T cell accumulation. To investigate whether this involved direct signalling of cDC1 by IL-4, we developed an *in vivo* model for cDC1 manipulation. Donor cDC1 were expanded *in vivo* in B6 mice^39^ prior to enrichment, CRISPR-Cas9-mediated deletion of either *Cd19* or *Il4ra* and then transfer into *Batf3*^−/-^ mice for replenishment of the cDC1 pool (**Figure 5c**). Two days later, PbT-I cells were transferred, and the mice vaccinated with RAS for assessment of PbT-I cell numbers 6 days later. Splenic PbT-I cell accumulation was impaired when cDC1 did not express IL-4Ra (sg*Il4ra*) (**Figure 5d**), suggesting that direct IL-4 signalling in cDC1 enhances the CD8 T cell response to RAS vaccination. Numbers of cDC1 recovered 6 days post-RAS vaccination were equivalent between groups (**Figure 5e and Supplementary Figure 5b**). Collectively, these experiments demonstrate that IL-4 acts directly on DC and on CD8 T cells for optimal CD8 T cell expansion in the context of RAS vaccination (**Figure 5f**).

### IL-4 synergises with CD40 to drive IL-12 production by cDC1

To identify the molecules induced by IL-4 in cDC1 that may contribute to the response to RAS vaccination, cDC1 gene expression was assessed *in vitro* following stimulation with IL-4 and αCD40. Expanded cDC1 were sort-purified and then cultured in each condition before isolation of RNA at 4 hours^40^. Sorted cDC1 were assessed after exposure to the following conditions: media alone, αCD40, IL-4, or αCD40+IL-4 (**Figure 6a**).

**Figure 6.**
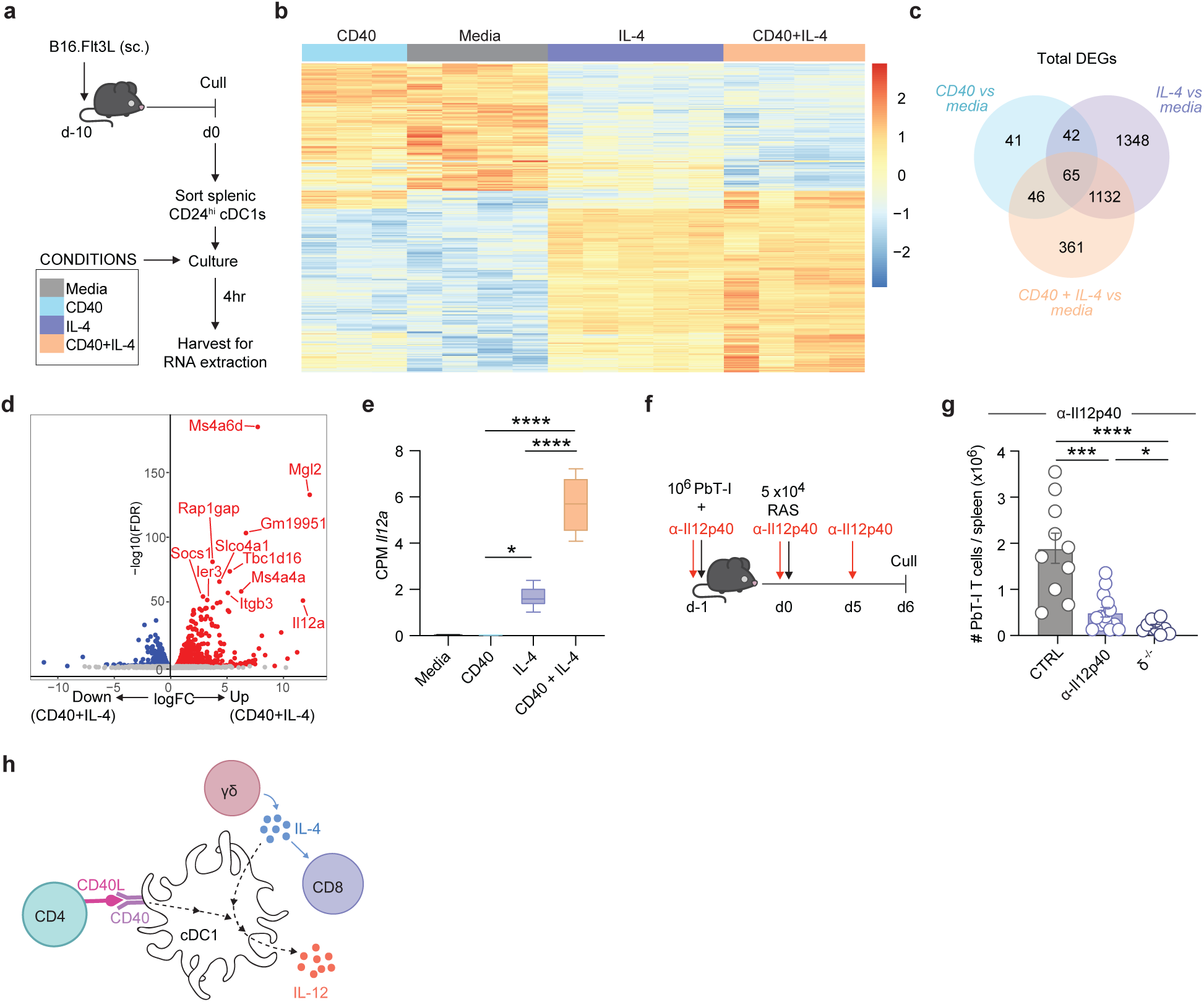
CD40 and IL-4 drive a unique gene expression profile in cDC1. **(a)** Experimental design for (**b-e**). **(b)** Heatmap showing the expression of differentially expressed genes (DEG) between media alone and αCD40+IL-4, across the different stimulation conditions. **(c)** Venn diagram showing DEGs between cDC1s cultured in media alone and cDC1s cultured with αCD40 only or IL-4 only or αCD40+IL-4 stimuli. **(d)** Volcano plot highlighting DEGs between αCD40 only and αCD40+IL-4 stimuli. **(e)** Gene expression of *Il12a* shown as counts per million (CPM). **(f)** Experimental design for (**g**). Mice received PbT-I cells one day prior to RAS vaccination and were treated with either α-IL-12p40 or an isotype control Ab on days –1, d0 and day 5. **(g)** Number of PbT-I cells in the spleen on day 6 post-RAS vaccination. **(h)** γδ T cell-derived IL-4 acts on cDC1s to promote IL-12 production which, along with IL-4, is required for CD8^+^ T cell accumulation. Data pooled from (**a-d**) n= 3-5 sequencing replicates derived from individual cDC1 cultures or (**f**) n=10 mice from 2 independent experiments. Error bars indicate mean + SEM. Data were log-transformed and compared using an ordinary one-way ANOVA. ****p<0.0001, ***p<0.001, **p<0.01, *p<0.05, ns=p>0.05.

Analysis of sequenced RNA revealed a marked transcriptomic shift in the presence of IL-4 that was augmented by the addition of αCD40 (**Figure 6b and c**). Specifically, there were 841 up-regulated and 762 down-regulated genes in response to αCD40+IL-4 when compared to media alone (**Figure 6b**). *Il12a*, was the most upregulated gene when comparing media with αCD40+IL-4 (LFC 11.17, FDR 1.5×10^−59^) and the second most upregulated gene when comparing αCD40+IL-4 with αCD40 alone (LFC 11.69, FDR 1.48×10^−52^) (**Figure 6d and e**). This gene encodes the p35 subunit of IL-12 which, when combined with the p40 subunit (which is constitutively expressed by cDC1), makes bioactive IL-12, a key stimulator of CD8 T cells. As it was only weakly upregulated in cDC1 exposed to IL-4 alone, efficient upregulation of *Il12a* also appeared dependent on the combination of both CD40 and IL-4 signals (**Figure 6e**).

To determine the contribution of bioactive IL-12 *in vivo* in the context of RAS vaccination, PbT-I cell accumulation was assessed after IL-12 blockade (**Figure 6f**). Assessment of PbT-I accumulation on day 6 showed significant impairment when IL-12 was blocked (**Figure 6g**). This supports a model where CD40 signalling synergises with IL-4 in cDC1 to induce IL-12 that is crucial for the accumulation of CD8 T cells in response to RAS vaccination (**Figure 6h**).

### IL-4 promotes CD8 T cell expansion by increasing IL12R expression in CD8 T cells

IL-4 and IL-12 were both essential for the enhanced accumulation of CD8 T cells in response to RAS vaccination. This led us to investigate whether these cytokines work synergistically to promote CD8 T cell expansion. To isolate the effects of the cytokines, we first examined the impact of IL-4 and IL-12 on CD8 T cell expansion *in vitro*.

PbT-I cells were activated by peptide coated antigen presenting cells in the presence of, no added cytokine, IL-4, IL-12, or both, and the number of cells retrieved from the cultures enumerated on day 4 and day 6. Limited cell growth was observed in the absence of exogenous cytokine (**Figure 7a**) and addition of either IL-4 or IL-12 only modestly impacted the number of cells recovered between day 4 and day 6 (**Figure 7b and c).** In contrast, addition of both IL-4 and IL-12 resulted in significantly higher cell recovery from the culture on day 6 (**Figure 7d**). These data suggested that the combination of IL-4 and IL-12 act synergistically in increasing the T cell response *in vitro*.

**Figure 7.**
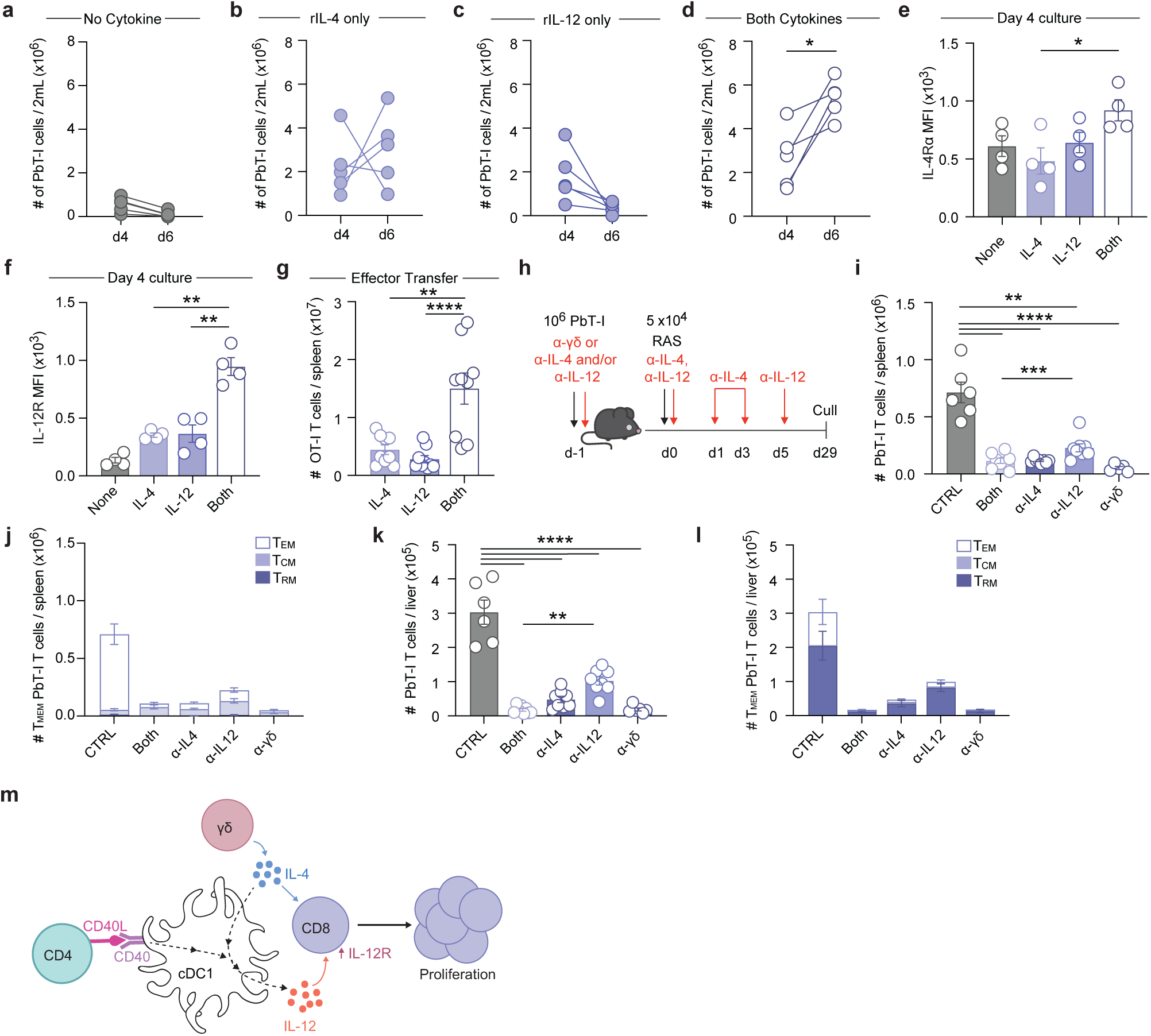
IL-4 and IL-12 synergise to promote expansion of CD8 T cells. PbT-I cells were peptide-activated *in vitro* in (**a**) media alone or in the presence of (**b**) 60 ng/mL rmIL-4, (**c**) 10 ng/mL rmIL-12 or (**d**) both rmIL-4 and rmIL-12. **(e)** IL-4R and (**f**) IL-12R surface expression on activated T cells 4 days after culture in the presence of different cytokines. OT-I T cells were activated *in vitro* in the presence of 60 ng/mL rmIL-4, 10 ng/mL rmIL-12 or both rmIL-4 and rmIL-12 then transferred into recipient mice at day 4 of culture. **(g)** Number of OT-I T cells in the spleen 3 days after effector transfer. **(h)** Experimental design for (**i-l**). Mice received 10^6^ PbT-I cells one day prior to RAS vaccination and were treated with α-IL-12p40 or α-IL-4 according to the indicated regimen. (**i and k**) Number of PbT-I cells in the (**i**) spleen and (**k**) liver, 29 days after vaccination. (**j and l**) Frequency of T_CM_, T_EM_ and T_RM_ within the CD44^+^ PbT-I cell compartment in the (**j**) spleen and (**l**) liver 29 days after RAS vaccination. (**m**) γδ T cell-derived IL-4 act on cDC1s to promote IL-12 production which, along with IL-4, signals directly on CD8^+^ T cells to drive proliferation and therefore enhance liver T_RM_ formation. Data pooled from (**a**) 5 (**e and f**) 4 (**g**) 3 or (**i-k**) 2 independent experiments where (**a-f**) has n = 4-5 individual cultures per group and (**g-l**) has n = 6-9 mice per group. Error bars indicate mean + SEM. Data were log-transformed then compared using (**a-d**) a paired t test or (**e-l**) an ordinary one-way ANOVA. ****p<0.0001, ***p<0.001, **p<0.01, *p<0.05, ns=p>0.05

To explore whether the synergy between IL-4 and IL-12 was mediated by changes in cytokine sensitivity resulting from alterations in receptor expression, we assessed IL-4R and IL-12R expression on T cells at day 4 of culture. Compared to addition of IL-4 alone, a combination of IL-4 and IL-12 induced a small increase in IL-4R (**Figure 7e**). More substantially, the addition of IL-4 + IL-12 increased expression of IL-12R over either cytokine alone (**Figure 7f**). These data suggested that the combination of IL-4 and IL-12 amplifies the sensitivity of CD8 T cells to IL-12 signalling through increased expression of IL-12R, thereby enhancing the CD8 T cell response. Further, *in vitro* activated OT-I T cells stimulated with IL-4 and IL-12 were maintained in greater numbers when transferred into mice (**Figure 7g**).

### γδ T cell-derived IL-4 and cDC1-derived IL-12 synergise to promote the generation of protective liver T_RM_ cells

Our data showed that in the absence of γδ T cells there is a defect in the initiation of the CD8 T cell response leading to a failure in the formation of T cell memory and defective generation of protective liver T_RM_ cells (**Figure 1e and f and Supplementary Figure 1f and g**). The important factor in this response was γδ T cell-derived IL-4, which induced cDC1 IL-12 production, and the two cytokines then synergised to promote CD8 T cell expansion. To confirm that removal of both IL-4 and IL-12 *in vivo* would recapitulate the lack of γδ T cells and therefore inhibit the formation of protective liver T_RM_ cells, we blocked IL-4 or IL-12, or both, in the context of RAS vaccination, and then examined T_RM_ numbers in the liver 29 days later (**Figure 7h**). Enumeration of the number of liver T_RM_ cells showed that IL-4 had a dominant effect on liver T_RM_ formation, but blockade of both cytokines phenocopied blockade of γδ T cells (with α-γδ Ab) in the spleen (**Figure 7i and j)** and in the liver (**Figure 7k and l**). These data showed that γδ T cells drive the response to RAS via delivery of IL-4 to DC, which in turn drives IL-12 production. The γδ T cell derived IL-4 and DC-derived IL-12 then synergise to enhance the expansion of CD8 T cells (**Figure 7m**), a proportion of which will differentiate into protective liver T_RM_ cells.

## Discussion

For full immunogenicity, DC must be effectively activated, either via strong PRR signalling or through CD4 T cell help via CD40L^1^. Here, we showed that the priming of CD8 T cells to *Plasmodium* sporozoite antigens depends on an additional layer of DC activation via IL-4, which is produced by γδ T cells. Here, IL-4 has a dual role: 1) when synergised with CD40 signals in DC, it promotes enhanced production of bioactive IL-12 and 2) by directly signalling CD8 T cells to increase IL-12R, it increases their sensitivity to IL-12 and thereby promotes expansion. In this vaccination setting, IL-4 is produced by Vγ1^+^ γδ T cells in the spleen, which become activated within the first 24 hours of injection of sporozoites.

The absolute requirement for γδ T cells for successful RAS vaccination in mice^20^ and the correlations with effective RAS vaccination in humans^20,29,41,42^ suggests that although *Plasmodium* sporozoites are reported to trigger toll like receptor 2 signalling^43^, this stimulation is not sufficient to push DC over a required activation threshold and that an additional IL-4 signal in combination with CD40 is required. Our findings likely extend beyond the context of RAS vaccination and may be applicable to other scenarios involving poor immunogens. In such cases, the sensing of infection or cellular damage by innate or innate-like cells may trigger IL-4 production, thereby enabling DC to effectively prime protective immune responses. One such innate-like T cell population are invariant natural killer T (NKT) cells, which are known to produce IL-4 in response to lipid ligands^44^. These ligands can be used as adjuvants to boost the response to RAS^45^ and to heat killed sporozoites^46^, likely via enhanced IL-4 production. However, the need for NKT cell-derived cytokines on top of CD4 T cell help in a four-cell paradigm, as we have shown here for γδ T cells, has not been described.

Here we identified that in mice, the Vγ1^+^ subset of γδ T cells are activated by RAS vaccination and these Vγ1^+^ cells provide a crucial function by supplying IL-4 for initiation of the CD8 T cell response. Previous studies demonstrated that provision of IL-4 by Vγ1^+^ γδ T cells had potent effects on B cells^47^ but Vγ1^+^ cells have not previously been directly linked to CD8 T cell responses *in vivo.* Despite identifying the specific subset involved and resolving the timing of activation, we have not yet successfully identified the trigger for Vγ1^+^ γδ T cell activation in response to RAS vaccination. Identification of the activation triggers for γδ T cells remain a challenge for the field^48,49^ but we hypothesise that the cells are responding to a parasite or parasite induced self-ligand exposed by the infectious process. In mice, γδ T cell-derived IL-17^50^ and IFN-γ^51^, induced by liver-stage parasites have been shown to play important roles in the pathology of *Plasmodium* blood stage infection. Here, we have shown that the initiation of the CD4 response to liver stage parasites is also dependent on a factor provided by γδ T cells but have not yet identified the nature of this factor. These concepts highlight that γδ T cells play a varied and substantial role in malaria disease.

Previous studies that described a role for γδ T cells in RAS vaccination proposed that a population of CD8 DC (now referred to as cDC1) that accumulated in the liver of RAS vaccinated mice when γδ T cells were present also correlated with the generation of hepatic CD8 T cell memory^20^. Although we have not directly investigated or ruled out a role for these hepatic DC in the memory component of the response to RAS, our data show that splenic cDC1 collaborate with γδ T cells to drive efficient priming of splenic CD8 T cells. It is possible that the inflammation induced by the combination of IL-4 and IL-12 observed when γδ T cells are present also leads to inflammation in the liver and the accumulation of CD8 cDC, commensurate with the accumulation of CD8 T cells in the liver. Given that the DC accumulation in the liver was observed late in the response (>60 days post-immunisation), this effect is unlikely to contribute to the crucial γδ T cell-dependent priming observed in the current study.

Prior to the discovery of T_RM_ cells, a role for IL-4 in RAS vaccination (for *P. yoelii* in BALB/c mice^16,17^) was identified. CD4 T cells, rather than γδ T cells, were implicated as the source of IL-4^16,17^. We speculate that a lack of CD40L signalling in the absence of CD4 T cells, rather than a lack of CD4 T cell-derived IL-4, was likely responsible for the phenotype observed. However, we currently cannot exclude the possibility that γδ T cell-derived IL-4 drives production of IL-4 by CD4 T cells. Nevertheless, we have shown that the initial and most important source of IL-4 is γδ T cell-derived.

Notably, we have shown that IL-4 and IL-12 are synergistic for CD8 T cell expansion due to enhanced expression of IL-12R, likely making the T cells more sensitive to bioactive IL-12. This is counterintuitive to published literature examining CD4 T cell responses, which states that IL-4-induced GATA3 activation suppresses IL-12 signalling^52^. However, synergistic effects of IL-4 and IL-12 have previously been reported for CD4 T cells^53^ and human T cells^54^, although the precise molecular mechanisms driving this synergy are currently unknown and require further investigation. Here, we show that IL-4 plays a key role in promoting the expansion of antigen-specific T cells not only by stimulating IL-12 production in DC but also enhancing CD8^+^ T cell sensitivity to the IL-12 produced. In the absence of IL-4 the entire response fails, making IL-4, and the triggers that induce its secretion by γδ T cells or other innate or innate-like cells, of major relevance to weakly stimulating pathogens (or tumours) that cannot intrinsically drive strong CD8 responses.

Importantly, a newly appreciated role for IL-4 in the context of immunotherapy has recently been highlighted. IL-4 production by B cells was shown to be one mechanism for successful PD-1 immunotherapy^55^ and two recent studies demonstrated the importance of an IL-4 driven type-2 response for effective and long-lived CAR T cell or anti-tumour responses^13,14^. These studies demonstrate that IL-4 has therapeutic implications beyond infectious diseases and indicate that γδ T cells should be investigated as a source of IL-4 in these models.

In conclusion, our study reveals that by providing IL-4, Vγ1^+^ γδ T cells drive the initiation of the response to a poorly immunogenic pathogen, transforming it into a robust trigger for CD8^+^ T cell immunity and tissue-tropic memory.

## Methods

### Mice, mosquitoes, and parasites

C57BL/6, B6.SJL-PtprcaPep3b/BoyJ (CD45.1), PbT-I GFP, PbT-II GFP, *Tcrd*^−/-56^, Trdc^tm1(EGFP/HBEGF/luc)Impr^ (TCRδ-GDL)^57^, *Il4*^−/-^, *Batf3*^−/-^, *Cd40l (Cd154)*^−/-^ , *Il17a*^−/-^ mice were bred and held at the Department of Microbiology and Immunology, The University of Melbourne, Australia. All experimental work and animal handling was conducted in strict accordance with the standards approved by the Animal Ethics Committee at the University of Melbourne (ethic project ID: 2015168, 20088). Experimental mice were sex matched and used at 6-16 wks of age. Both male and female mice were used. TCRδ-GDL^57^ mice were kindly provided by I. Prinz (University Medical Center Hamburg-Eppendorf).

*Plasmodium berghei* ANKA (PbA) wild-type Cl15cy1 (BEI Resources, NIAID, NIH: MRA-871) or *P. berghei* CS5M, which express the cognate antigen for OT-I T cells^58^, parasites were used. Sporozoites were generated in 4-5-week old male Swiss Webster mice purchased from the Monash Animal Service and held at the School of Botany, The University of Melbourne, Australia. *Anopheles stephensi* mosquitoes (strain STE2/MRA-128 from BEI Resources, The Malaria Research and Reference Reagent Resource Centre) were reared and infected as described previously^59^. Sporozoites were hand-dissected from mosquito salivary glands, resuspended in cold PBS then irradiated with 20 K rads from a gamma ^60^Co source (WEHI irradiation facility). Irradiated sporozoites were diluted to a total number of 5 ×10^4^ per mouse and injected via the tail vein using 26G insulin syringes.

In blood-stage malaria experiments, donor mice were injected with frozen stabilates of blood-stage P.*berghei* ANKA (PbA) wild-type Cl15cy1 (BEI Resources, NIAID, NIH: MRA-871). Three to seven days later, mice were bled for parasitemia, the RBCs irradiated with 20 K rads from a gamma ^60^Co source (WEHI irradiation facility), and recipient mice were injected with 5 ×10^4^ irradiated PbA-infected RBCs diluted in 0.2 mL of sterile PBS.

### Cell Enrichments

#### T cells

Naïve CD8^+^ PbT-I or CD4^+^ PbT-II cells were negatively enriched from the spleens and lymph nodes of PbT-I/uGFP, OT-I/uGFP or PbT-II/uGFP transgenic mice. Briefly, cells were passed through a 70 µM filter sieve to obtain a single-cell suspension and red blood cells lysed. Single-cell suspensions were labelled with rat monoclonal antibodies specific for MHC Class II (M5-114), Mac-1 (M1/70, F4/80), Gr-1 (RB6-8C5) and either CD4 (GK1.4) (for PbT-I and OT-I CD8 T cells) or CD8 (53-6.7) (for PbT-II CD4 T cells) prior to incubation with BioMag goat anti-rat IgG coupled magnetic beads (Qiagen) and magnetic separation. Enriched T cells were typically 80-95% pure. 1 ×10^6^ purified T cells in 0.2 mL of sterile PBS were injected via the tail vein into recipient mice. Where co-transfers were performed, a total number of 2 ×10^6^ purified T cells were injected via the tail vein where cell types were transferred at a ratio of 1:1.

#### γδ T cells

Naïve γδ T cells were negatively enriched from spleens of B6 or *Il4*^−/-^ mice. Single cell-suspensions were achieved by finely mincing and digesting spleens in 1 mg/mL Collagenase Type 3, 20 µg/mL DNase I supplemented with a competitive antagonist of P2X_7_R (A-438079 hydrochloride; Santa Cruz Biotechnology) with intermittent agitation. Undigested fragments were removed by filtering through a 70 µM mesh followed by red blood cell lysis. Single cell suspensions were labelled with rat monoclonal antibodies specific for MHC Class II (M5-114), Mac-1 (M1/70, F4/80), CD4 (GK1.4), CD8 (53-6.7) and B220 (RA3-6B2) prior to incubation with BioMag goat anti-rat IgG coupled magnetic beads (Qiagen) and magnetic separation. 2×10^6^ enriched γδ T cells in 0.2 mL of sterile PBS were injected via the tail vein into recipient mice.

#### Dendritic cells

Dendritic cells were isolated from spleens of B16.Flt3L tumour-bearing mice. Briefly, spleens were finely minced in 1 mg/ml Collagenase Type 3, 20 µg/ml DNase I and continuously agitated for 15 minutes. DC-T cell complexes were disrupted with the addition of 0.1M EDTA (pH7.2). Undigested fragments were removed by filtering through a 70 µM mesh before light-density separation with 1.077g/cm^3^ Nycodenz medium via centrifugation at 1700rpm at 4°C for 12min. To increase the purity of cDC1 precursors, the light density fraction was collected, and cDC1s/pre-cDC1s were negatively enriched for through incubation with rat mAb against CD3ε (KT3), Thy1.1 (T24/31.7), Gr-1 (RB6-8C5), B220 (RA3-6B2), Ter-119 and CD11b (M1/70) prior to incubation with BioMag goat anti-rat IgG coupled magnetic beads (Qiagen) and magnetic separation. Target cells were characterised as CD11c^+^, MHC-II^int/+^, CD11b^−^, CD24^hi^, CD8α^+^ with purities ranging between 60-80%. 2-5 × 10^6^ purified cDC1s in 0.2 mL of sterile PBS were injected via the tail vein into recipient mice, 2 days prior to PbT-I cell transfer.

### CRISPR/Cas9 gene-editing

#### T cells

Purified CD8^+^ PbT-I or CD4^+^ PbT-II cells were gene-edited via electroporation with single guide (sg) RNA ribonucleoproteins (RNPs) (sgRNA/Cas9 RNPs). Briefly, sgRNA/Cas9 RNPs were formed in Nuclease-free H_2_O using 0.6 µl of Alt-R S.*pyogenes* Cas9 Nuclease V3 (10 mg/mL, Integrated DNA Technologies) incubated with 0.3 nmol of sgRNA targeting either *Il4r*α (5’-AGUGGAGUCCUAGCAUCACG-3’, 3’-AUCCAGGAACCACUCACACG-5’) or *Cd19* (5’-AAUGUCUCAGACCAUAUGGG-3’) purchased from Synthego (CRISPRevolution sgRNA EZ Kit) for 10 mins at room temperature. For each sgRNA/Cas9 electroporation reaction, 10 × 10^6^ target cells were required. In each reaction, target cells were resuspended in 20 µl of reconstituted P3 buffer then mixed with the sgRNA/Cas9 RNP complex and electroporated using a Lonza 4D-Nucleofector (program code: DN100). Electroporated cells were then rested in a 96-well plate for at least 10 minutes at 37°C. For injection into mice, 10^6^ cells were injected into mice via the tail vein in 0.2 mL of PBS, one day prior to RAS vaccination.

#### Dendritic Cells

Purified pre-cDC1s were gene-edited via electroporation with single guide (sg) RNA ribonucleoproteins (RNPs) (sgRNA/Cas9 RNPs) using an optimised protocol. sgRNA/Cas9 RNPs were formed as described for T cell gene-editing. For each sgRNA/Cas9 electroporation reaction, 10 × 10^6^ target cells were required. In each reaction, target cells were resuspended in 20 µl of reconstituted P3 buffer then mixed with the sgRNA/Cas9 RNP complex and electroporated using a Lonza 4D-Nucleofector (program code: CM137). Electroporated cells were then rested in a 96-well plate for at least 20 minutes at 37°C then injected intravenously into *Batf3*^−/-^ mice three days prior to immunisation. All mice received PbT-I cells one day prior to RAS immunisation.

#### *In vitro* cDC1 gene-expression assay

Dendritic cells were isolated through the protocol outlined above. CD11c^+^, MHC-II^int/+^, CD11b^−^, CD24^hi^, F4/80^−^, B220^−^, CD172α^−^ cells were then sorted using a BD FACSAria^TM^ III Cell Sorter into filtered RPMI 1640 supplemented with 50% heat-inactivated FCS and pelleted. In a 24-well plate, 5 × 10^5^ cells were plated in complete Kenneth D Shortman RPMI (KDS-RPMI 1640, 10% FCS, 2 mM L-glutamine, 100 U/ml penicillin, 100 mg/mL streptomycin and 50 mM 2-mercaptoethanol). To assess cDC1 gene expression in response to IL-4, cells were incubated in the presence of α-CD40 (10 μg/ml), recombinant mouse IL-4 (60 ng/mL), IFN-γ (20 ng/mL), or LPS (1 μg/ml). Plates were then incubated at 37°C and 5% CO_2_ for 4 hours. For RNA isolation, cells were washed and resuspended using TRIzol (Life Technologies), snap frozen and stored at -80°C.

### RNA Isolation and sequencing

To determine gene expression in cDC1s, mRNA was extracted using a Direct-zol RNA MicroPrep kit (Zymo Research) as per manufacturer’s protocol. Libraries were prepared using Illumina stranded mRNA library kits and the outputs sequenced to a depth of 20M reads per sample on a 150PE on Illumina NovaSeq X Plus 10B flow cell. Library preparation and sequencing was conducted at the Australian Genome Research Facility.

### Bioinformatics Analyses

Sequencing reads were aligned to the GRCm39 reference genome and transcriptome (version 105) using the STAR aligner (version 2.7.8a)^60^ and transcript counts established using featureCounts from the subread package (version 2.0.0)^61^. The data was then analysed using R (version 4.2.0), limma (version 4.54.1)^62^ and plots generated with ggplot2^63^. We first merged technical replicates from separate sequencing lanes into their respective biological replicates by summing up the counts. To obtain differentially expressed genes between the experimental groups, the data were filtered for lowly or non-expressed genes (filterByExpr function) and normalised counts by Trimmed Mean of M-values (TMM). Mean-variance trends were estimated with the voom function, and a linear model was fit using lmFit. The model parameters were moderated using eBayes, p values corrected for multiple testing using the Benjamini-Hochberg method and differential genes obtained for each of the pairwise contrasts between the experimental groups. The heatmap of expression values of top differential genes was generated with the pheatmap package (version 1.0.12) on log2 normalised CPM values.

### Bone-marrow chimeras

Mice were irradiated with two doses of 550 rad 3 hours apart. Mice were reconstituted with 3-5 × 10^6^ bone marrow (BM) cells from either B6, *Il4*^−/-^ or *Tcrd*^−/-^ mice. In the case of mixed bone-marrow chimeras, mice were reconstituted with a 1:1 ratio of bone-marrow cells from B6, *Il4*^−/-^ and/or *Tcrd*^−/-^ mice totalling to 3-5 × 10^6^ BM cells per mouse. BM cells were prepared by removing the tibia and femur of donor mice. Bones were flushed with complete RPMI using a 23G needle and a 10 mL syringe then passed through a 70 µM mesh filter to collect bone marrow. T cells were removed by resuspending the pellet with Abs against CD4 (RL172), CD8α (3.168), and Thy1.1 (Jlj) for 30 minutes at 4°C followed by incubation with rabbit complement for 20 minutes at 37°C to remove bound cells. All irradiated mice were injected via the tail vein with 3-5 × 10^6^ bone marrow cells in 0.2 mL of PBS. The following day, chimeras were injected intraperitoneally with 0.1 mL of T24 (α-Thy1.1). Irradiated mice received antibiotic water containing 2.5 g/L Neomycin Sulphate and 0.94 g/L Polymyxin B sulphate for the first 4 weeks after irradiation followed by normal water for at least another 4 weeks prior to experimental work.

#### *In vivo* antibody depletion/blockade

Mice were treated intraperitoneally with purified α-IL-4 (clone 11B11) for 6 days starting one day prior to RAS vaccination. For the first 3 days, mice were treated with 500 µg then 200 µg for the final 3 days. Control mice were treated with α-horseradish peroxidase (clone HRPN) using the same dose and regimen^16^.

For IL-12p40 neutralisation, mice were treated intraperitoneally with 250 µg of purified α-IL-12p40 (clone C17.8) for 2 days beginning one day prior to RAS vaccination with an additional dose the day prior to the end point. Control mice were treated with α-trinitrophenol (clone 2A3) at the same dose and schedule^64^.

To block γδ T cell function *in vivo*, mice received a single 200 µg dose of α-pan-γδτCR (clone GL3) or α-Vγ1 (clone 2.11)^65^ intravenously on the day prior to immunisation with RAS. As a control, separate mice were treated with 200 µg of Purified Armenian Hamster IgG (clone HTK888)^20^.

#### Organ processing and flow cytometry

Single cell suspensions were prepared from spleen or livers by passing the tissues through 70 µM mesh filters. For experiments in which splenic DCs were enumerated, spleens were finely minced in 1mg/ml Collagenase Type 3, 20 µg/ml DNase I and continuously agitated for 15 minutes. DC-T cell complexes were disrupted with the addition of 0.1M EDTA (pH7.2) prior to filtration through a 70 µM mesh. Liver preparation required an additional step of resuspending cells in 35% isotonic Percoll and centrifugation at 500 rpm for 20 minutes at room temperature. Red blood cells were removed from spleen and liver pellets and labelled with a near-IR fixable live-dead dye to exclude dead cells. Fluorochrome-conjugated monoclonal antibodies specific for TCRδ, CD3ε, NK1.1, CD8α, CD4, B220, IL-4R, TCRβ, CD69, CD62L, CD44, CD27, Vγ1, Vγ4, CD11a, CXCR6, CD11c, CD11b, XCR1, CD24, CD172a and MHC-II were used for surface staining and performed for an hour at 4°C. H2-K^b^-PbRPL6_120-127,_ tetramers were provided by Professor Stephanie Gras. Sphero Blank Calibration beads were used to generate cell counts. Cells were acquired on a Cytek Aurora using the SpectroFlo v3.0 system (Cytek) and further analysed on FlowJo (TreeStar, Inc).

#### *In-vitro* activation and culture of T cells

PbT-I cells were activated in complete RPMI (RPMI 1640, 10% FCS, 2mM L-glutamine, 100 U/mL penicillin, 100 mg/mL streptomycin and 50 mM 2-mercaptoethanol) for up to 6 days with PbRPL6_120-127_ (NVFDFNNL) peptide-pulsed splenocytes and in the presence of LPS (1 μg/mL; Sigma-Aldrich) and either: no added cytokine, recombinant mouse IL-12 (10 ng/mL^66^), recombinant mouse IL-4 (60 ng/mL^34^; BioLegend) or both IL-12 and IL-4. Activated cells were fed with complete RPMI on day 3 and split on day 4. To assess phenotype, cells were washed and labelled with a near-IR fixable live-dead dye to exclude dead cells and fluorochrome-conjugated monoclonal antibodies specific for TCRβ, IL-4R, IL-12R, CD44, Vα8.3, and CD8α.

### Statistical Analyses

Statistical analyses of graphed data were performed using GraphPad Prism software. Data are shown as mean values ± SEM. The statistical tests used, and p values are indicated in each figure legend. All statistical tests were performed on log-transformed data unless otherwise specified.

## Acknowledgments

The work was supported by The Australian National Health and Medical Research Council (Grant #1105817 and 2002682) and The Australian Research Council (Grant #DP220103545). S.G. is supported by an NHMRC SRF fellowship (#1159272). We wish to acknowledge the University of Melbourne Bioresources platform and the Melbourne Cytometry platform for support and expertise. We also wish to thank Melanie Damtsis and Gayle Davey for all the indirect contributions that make great research possible and Dr Yannick Alexandre for critical reading of the manuscript.

## Author Contributions

SL performed the experiments with help from DM, SLiu, ZG, RM and LB and wrote the draft of the paper. AC provided RAS and CJ both under supervision by GM. CX and TB contributed methodology supervised by FK or LM. SG and IP provided resources. JS conducted bioinformatic analysis of the RNAseq data. AB, SB, IC and DFR helped with conceptualization and methodology. WRH and LB conceptualized, funded and supervised the study and wrote the paper.

