## supplementary figures for "Vγ1^+^ γδ T cell-derived IL-4 initiates CD8 T cell immunity"

#### Supplementary Figure 1

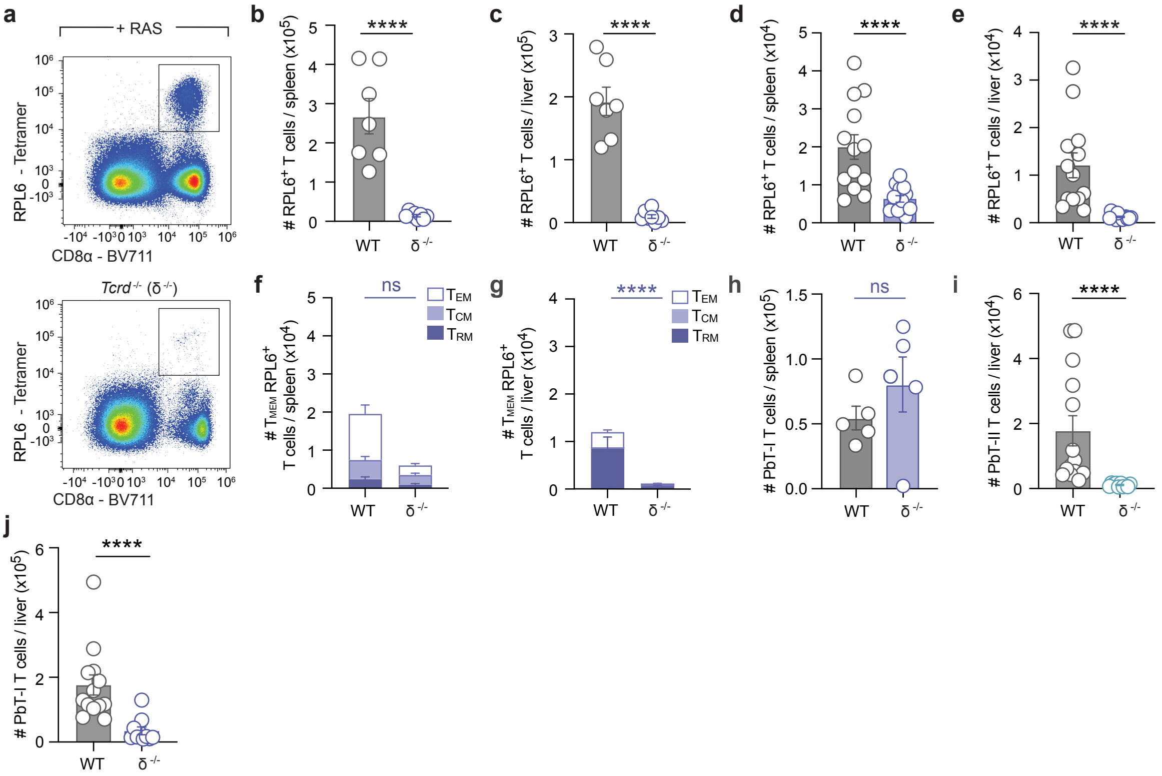

#### Supplementary Figure 2

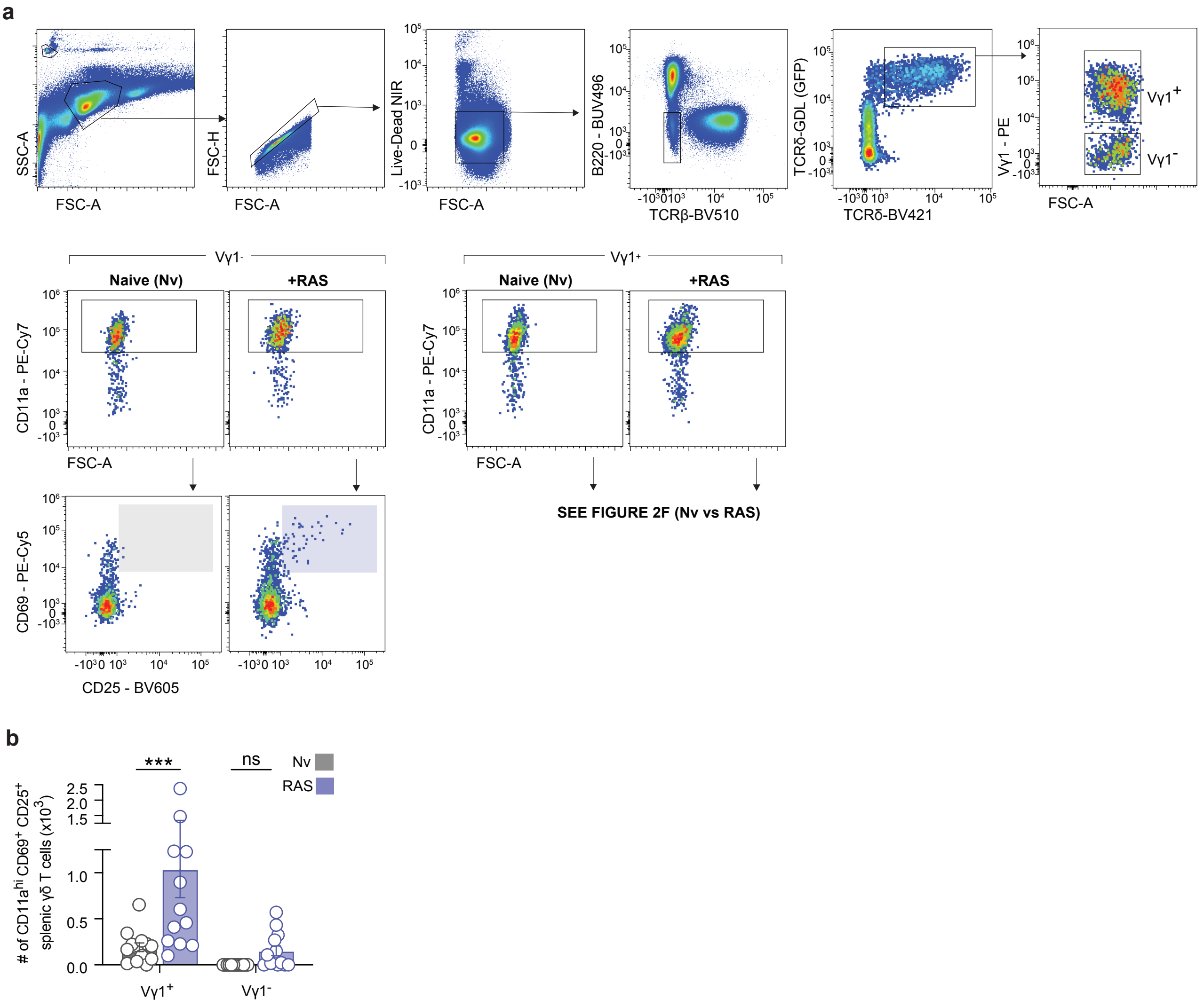

#### Supplementary Figure 3

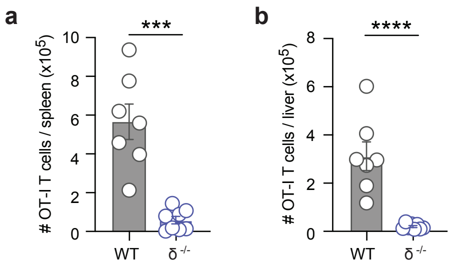

#### Supplementary Figure 4

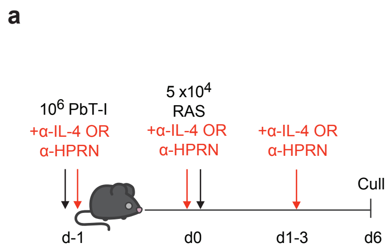


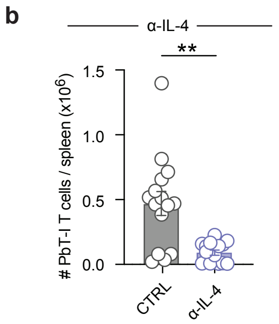

### Supplementary Figure 5

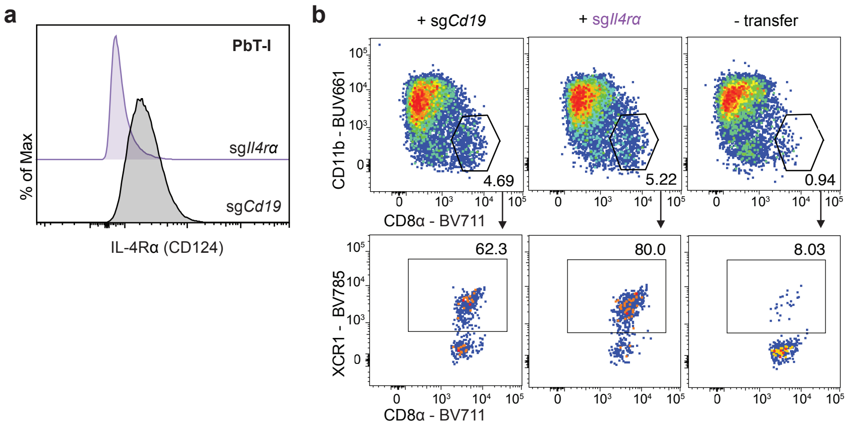
